# Anti-Viral and Anti-Inflammatory Therapeutic Effect of RAGE-Ig Protein Against Multiple SARS-CoV-2 Variants of Concern Demonstrated in K18-hACE2 Mouse and Syrian Golden Hamster Models

**DOI:** 10.1101/2023.06.07.544133

**Authors:** Nisha Rajeswari Dhanushkodi, Swayam Prakash, Afshana Quadiri, Latifa Zayou, Ruchi Srivastava, Amin Mohammed Shaik, Berfin Suzer, Izabela Coimbra Ibraim, Gary Landucci, Delia F Tifrea, Mahmoud Singer, Leila Jamal, Robert A Edwards, Hawa Vahed, Lawrence Brown, Lbachir BenMohamed

## Abstract

**Significance:** SARS-CoV-2 Variants of Concern (VOCs) continue to evolve and re-emerge with chronic inflammatory long-COVID sequelae necessitating the development of anti-inflammatory therapeutic molecules. Therapeutic effects of the Receptor for Advanced Glycation End products (RAGE) were reported in many inflammatory diseases. However, a therapeutic effect of the RAGE in COVID-19 has not been reported. In the present study, we investigated whether and how the RAGE-Ig fusion protein would have an anti-viral and anti-inflammatory therapeutic effect in the COVID-19 system.

**Methods:** The protective therapeutic effect of RAGE-Ig was determined in vitro in K18-hACE2 transgenic mice and Syrian golden hamsters infected with six various VOCs of SARS-CoV-2. The underlying anti-viral mechanism of RAGE-Ig was determined *in vitro* in SARS-CoV-2-infected human lung epithelial cells (BEAS-2B).

**Results:** Following treatment of K18-hACE2 mice and hamsters infected with various SARS-CoV-2 VOCs with RAGE-Ig, we demonstrated: (*i*) significant dose-dependent protection (i.e. greater survival, less weight loss, lower virus replication in the lungs); (*ii*) a reduction of inflammatory macrophages (F4/80^+^/Ly6C^+^) and neutrophils (CD11b^+^/Ly6G^+^) infiltrating the infected lungs; (*iii*) a RAGE-Ig dose-dependent increase in the expression of type I interferons (IFN-α, and IFN-β) and type III interferon (IFNλ_2_) and a decrease in the inflammatory cytokines (IL-6 and IL-8) in SARS-CoV-2-infected human lung epithelial cells; and (*iv*) a dose-dependent decrease in the expression of CD64 (FcgR1) on monocytes and lung epithelial cells from symptomatic COVID-19 patients.

**Conclusion:** Our pre-clinical findings revealed type I and III interferons-mediated anti-viral and anti-inflammatory therapeutic effects of RAGE-Ig protein against COVID-19 caused by multiple SARS-CoV-2 VOCs.

## INTRODUCTION

The Coronavirus disease 2019 (COVID-19) pandemic has created one of the largest global health crises in almost a century. Although the current rate of SARS-CoV-2 infections has decreased significantly; the long-term outlook of COVID-19 remains a serious cause of high death worldwide; with the mortality rate still surpassing even the worst mortality rates recorded for the influenza viruses. The continuous emergence of SARS-CoV-2 variants of concern (VOCs), including multiple heavily mutated Omicron sub-variants, have prolonged the COVID-19 pandemic and outlines the urgent need for a next-generation vaccine that will protect from multiple SARS-CoV-2 VOCs. COVID-19 patients are reported with various pulmonary ailments known to be associated with the accumulation of inflammatory molecules like advanced glycation end-products (AGEs), calprotectin, high mobility group box family-1 (HMGB1), cytokines, angiotensin-converting enzyme 2 (ACE2), and other molecules in the alveolar space of lungs as well as plasma (1–6). The receptor for Advanced Glycation End Products (RAGE), is the AGEs-specific multi-ligand receptor involved in inflammatory responses to various chronic inflammatory lung diseases (7, 8). RAGE is known to be highly expressed in the lungs, especially in the alveolar epithelial cells (AECs), in comparison to other organs of the human body. The RAGE pathway has been reported in the pathogenesis of lung diseases such as chronic obstructive pulmonary disease (COPD), interstitial lung diseases, and ARDS (9, 10). In recent years, the therapeutic intervention of the RAGE pathway has been demonstrated to benefit patients suffering from ARDS, COPD, pulmonary fibrosis, and other pathologies associated with the pulmonary system (11–15). Notably, enhanced levels of RAGE ligands are associated with inflammatory disorders Fields(16–18), such as diabetes or other chronic inflammatory disorders. This receptor has a causative effect on a range of inflammatory diseases.

In the context of COVID-19, the RAGE axis is observed to be associated with comorbidities such as COPD, coronary artery disease (CAD), atherosclerosis, and hypertension (19–29). Patients with such co-morbidities are at an increased risk of exacerbation and decompensation when infected with SARS-CoV-2, given the severe inflammatory response generated. Interestingly, Broncho alveolar lavage (BAL) and serum from COVID-19 patients have detectable soluble RAGE protein that might potentially have a prognostic role. In COVID-19 patients, S100A12-like RAGE ligands are considered potential biomarkers that correlate with the severity of lung inflammation (2, 30). Although recent studies have explored the therapeutic interventional strategies for the RAGE pathway in chronic inflammatory diseases, strategies that modulate the RAGE pathway have yet to be studied extensively in the realm of the COVID-19 (31, 32).

The RAGE receptor exists either as membrane-bound RAGE (mRAGE) or a soluble form (sRAGE) (33). Ligands such as AGEs, S100/calgranulin, and HMGB1 protein initiate intracellular signal stimulation when linked to mRAGE (2). RAGE/AGE axis stimulation causes the downstream activation of various inflammatory transcription factors, including nuclear factor kappa B (NFkB) (34, 35). sRAGE can prevent ligands from binding to mRAGE, thereby inhibiting inflammatory activation (8). Isoforms of the RAGE protein, which lack the transmembrane and the signaling domain (commonly referred to as sRAGE) provide a means to develop a cure against RAGE-associated diseases.

In this report, we used a RAGE fusion protein that consists of a RAGE ligand-binding element, a heavy-chain immunoglobulin of a G4 isotype constant domain, and a linker connecting the ligand-binding element with the constant domain. This RAGE-Fc (fragment, crystallizable region of the antibody) binds to ligands of RAGE through competition and inhibits RAGE signaling by competing with membrane-bound receptors for binding ligands (hereafter referred to as RAGE-Ig). We subsequently evaluated how efficiently RAGE-Ig protein may be used to treat COVID-19 in the context of multiple SARS-CoV-2 variants of concerns (VOCs), for which no effective treatment is available (*2*). One recent study reported the effect of RAGE antagonists to be effective against COVID-19 in the mouse model (*36*). To our knowledge, we are the first group to test the pre-clinical therapeutic effect of this RAGE-Ig fusion protein against SARS-CoV-2 VOCs infection in both mouse and hamster models. Our findings from this report also demonstrate a unique coupled anti-viral and anti-inflammatory effect of the RAGE-Ig along with their respective mechanisms involved.

## MATERIALS AND METHODS

### Viruses

SARS-CoV-2 viruses specific to four variants, namely (i) SARS-CoV-2-USA/WA/2020 (Batch Number: G2027B); (ii) Alpha (B.1.1.7) (isolate England/204820464/2020 Batch Number: C2108K); (iii) Beta (B.1.351) (isolate South Africa/KRISP-EC-K005321/2020; Batch Number: C2108F), and (iv) Gamma (P.1) (isolate hCoV-19/Japan/TY7-503/2021; Batch Number: G2126A) were procured from Microbiologics (St. Cloud, MN). The initial batches of viral stocks were propagated to generate high-titer virus stocks. Vero E6 (ATCC-CRL1586) cells were used for this purpose using an earlier published protocol (37). Procedures were completed only after appropriate safety training was obtained using an aseptic technique under BSL-3 containment.

### Animal Infection and RAGE-Ig treatment

The University of California-Irvine (Irvine, CA) conformed to the Guide for the Care and Use of Laboratory Animals published by the US National Institute of Health (IACUC protocol #22-086). Male and female K18-hACE2 transgenic mice were intranasally infected with 1X10^4^ pfu of SARS-CoV-2 (USA-WA1/2020) in 20ul and treated with 100 μg RAGE-Ig/mouse by intraperitoneal (i.p.) or subcutaneous (s.c.) or mock-treated on alternate days (n = 10 each group) with 100 μg of RAGE per mouse. The RAGE-Ig treatment was given on alternate days (beginning from day 1 to day 9 post-infection (p.i.). Mice were monitored daily for weight loss until day 14 p.i. For the dose-kinetics experiment, mice were treated with different doses of RAGE-Ig (100, 50, 25 μg) per mouse on alternate days (beginning from day 1 to day 9). Briefly, K18-hACE2 transgenic mice were intranasally infected with 1X10^4^ pfu of SARS-CoV-2 (USA-WA1/2020) in 20ul and treated with RAGE-Ig (s.c.) at 100 or 50, or 25 ug/mouse/dose (n = 4) each on alternate days from day1 to day 9 p.i. (1,3,5,7&9 days p.i) as indicated. At day 10 p.i, mice were euthanized and immune cells from the lung were used for flow cytometry. For the hamster experiments, golden Syrian hamsters (7-8 weeks old) (n = 20) were infected with SARS-CoV-2 WA (3 x 10^5^ pfu/hamster) or Delta (6.9X10^4^ pfu/hamster). mRAGE treatment (1.5 or 3mg/hamster) was administered by subcutaneous injection of RAGE-Ig or vehicle control on alternate days from day 1 to day 9 p.i and the animals were monitored for weight loss every day until day 14 p.i.

### Flow cytometry

Single-cell suspensions from the mouse lungs after collagenase treatment (10 mg/ml) for 1 hour were used for FACS staining. The following antibodies were used: anti-mouse CD3 (clone 17A-2, BD Biosciences), CD11c (clone HL3, BD Biosciences), CD11b (clone M1/70, BD Biosciences), F4/80 (clone BM8, BD Biosciences), F4/80, Ly6G, Ly6C, CD64, HLA-DR (clone M5/114, BD Biosciences). For surface staining, mAbs for various cell markers were added to a total of 1 x10^6^ cells in phosphate-buffered saline containing 1% FBS and 0.1% Sodium azide (fluorescence-activated cell sorter [FACS] buffer) and left for 45 minutes at 4°C. For intracellular/intranuclear staining, cells were first treated with cytofix/cytoplasm (BD Biosciences) for 30 minutes. Upon washing with Perm/Wash buffer, mAbs were added to the cells and incubated for 45 minutes on ice and in the dark. Cells were washed again with Perm/TF Wash and FACS buffer and fixed in PBS containing 2% paraformaldehyde (Sigma-Aldrich, St. Louis, MO). A total of 100,000 events were acquired by the LSRII (Becton Dickinson, Mountain View, CA) followed by analysis using FlowJo software (TreeStar, Ashland, OR).

### Histology of animal lung

Mouse lungs were preserved in 10% neutral buffered formalin for 48hr before transferring to 70 % ethanol. The tissue sections were then embedded in paraffin blocks and sectioned at 8 μm thickness. Slides were deparaffinized and rehydrated before staining for hematoxylin and eosin for routine immunopathology.

### Viral PCR assay

Lung tissue was analyzed for SARS-CoV-2 specific RNA by qRT-PCR. As recommended by the CDC, we used ORF1ab-specific primers (Forward-5’-CCCTGTGGGTTTTACACTTAA-3’ and Reverse-5’-ACGATTGTGCATCAGCTGA-3’) and probe (6FAM-CCGTCTGCGGTATGTGGAAAGGTTATGG-BHQ) to detect the viral RNA level in lung.

### PBMC isolation

Peripheral blood was collected and PBMCs (peripheral blood mononuclear cells) were isolated by Ficoll-opaque density gradient. The cells were washed in PBS and re-suspended in a complete culture medium consisting of RPMI-1640 medium containing 10% FBS (Bio-Products, Woodland, CA) supplemented with 1x penicillin/L-glutamine/streptomycin, 1x sodium pyruvate, 1x non-essential amino acids. About 1 million PBMCs were treated with different concentrations of RAGE-Ig, and the supernatant was used for cytokine ELISA while the cells were used for flow cytometry. Human CD14^+^ monocytes were isolated from PBMCs and further differentiated into uncommitted macrophages (M0) by macrophage colony-stimulating factor (M-CSF). The effect of RAGE-Ig on the polarization of M0 into classically activated (M1) or activated (M2) macrophages (using a combination of IFN-γ +/-RAGE-Ig for M1 or IL-4 +/-RAGE-Ig for M2) was studied. The RAGE-Ig protein treatment causes dose-dependent morphological changes in M0 macrophages during monocyte differentiation into macrophages.

### Invitro assays using cell line

Human lung epithelial cells BEAS-2B cells (CRL-9609, ATCC) were used for in vitro assays. Briefly, 1 million cells were treated with different concentrations of RAGE-Ig, and the supernatant was used for cytokine ELISA. ELISA or Luminex for various cytokines such as IFN-γ, IL-8, IL-6, IFN-Β, IFN-Λ3 was performed as per manufacturer’s (PBL Assay science, Millipore) instructions.

### Statistical analysis

Data for each assay were compared by ANOVA and Student’s t-test using GraphPad Prism version 5 (La Jolla, CA). Differences between the groups were identified by ANOVA and multiple comparison procedures. Data are expressed as the mean ± SD. Results were considered statistically significant at a P value of ≤ 0.05.

### RAGE-Ig Protein

The RAGE-Ig protein used in these studies was manufactured by Catalent Biologics (Madison, WI) and provided by Galactica Pharmaceuticals, Inc. (Villanova, PA).

## RESULTS

### 1. Dose-dependent protection in RAGE-Ig-treated SARS-CoV-2-infected K18-hACE2 mice is associated with a decrease in lung inflammation

RAGE inhibitors may be used as novel therapeutic targets for prevention, regression, and slowing of the progression of SARS-CoV-2 infections. To examine the effect of RAGE-Ig treatment in SARS-CoV-2 infection, the best route of drug administration was studied. Mice were treated with the RAGE-Ig by various routes of treatment (either i.p. or s.c.) after infection with SARS-CoV-2. Eight-to nine-week-old male and female K18-hACE2 transgenic mice were intranasally infected with 1 x 10^4^ pfu of SARS-CoV-2 (USA-WA1/2020) in 20 μl treated with 100 μg RAGE-Ig/mouse by either the intraperitoneal (i.p.) (*n* = 10) or subcutaneous (s.c.) route (*n* = 10) or mock-treated (*n* = 10) on alternate days (1, 3, 5, 7 and 9 days p.i.) (**Fig. 1A**). Mice that were treated subcutaneously with100 μg RAGE-Ig were found to be significantly protected from weight loss and mortality (**Fig. 1B**). Next, we studied the dose-response of RAGE-Ig fusion protein treatment on disease outcomes in SARS-CoV-2 infection. Eight-to nine-week-old male and female K18-hACE2 transgenic mice were intranasally infected with 1 x 10^4^ pfu of SARS-CoV-2 (USA-WA1/2020) and treated with RAGE-Ig (s.c.) at 100μg/mouse/dose (*n* = 4) or 50μg/mouse/dose (*n* = 4) or 25μg/mouse/dose (*n* = 4) or mock-treated (*n* = 4) on alternate days (1, 3, 5, 7 and 9 days p.i.) (**Fig. 1C**). We found a dose-dependent survival and weight loss benefit upon treatment with RAGE-Ig (**Fig. 1D**). The 100 μg dose was found to be the most effective dose that protected against severe symptoms of COVID-19 in mice. Additionally, a significant dose-dependent decrease in inflammatory cells found in the lung interstitium of mice treated with RAGE-Ig after SARS-CoV-2 infection was detected; notably with macrophages (Ly6C^+^F4/80^+^MHCII^+^) and the monocytes (Ly6C^+^F4/80^+^MHCII^-^) in the lungs (**Supplemental Fig. S1** and **Fig. 1E**), interstitial macrophages (CD11b^+^CD64^+^CD45^+^) and neutrophils (CD11b^+^LY6G^+^CD45^+^). These cells are associated with the inflammatory response in the lungs of mice infected with SARS-CoV-2. Altogether, this report suggests the potential therapeutic use of RAGE-Ig protein to attenuate macrophage-related inflammatory reactions in the treatment of COVID-19.

**Figure 1.**
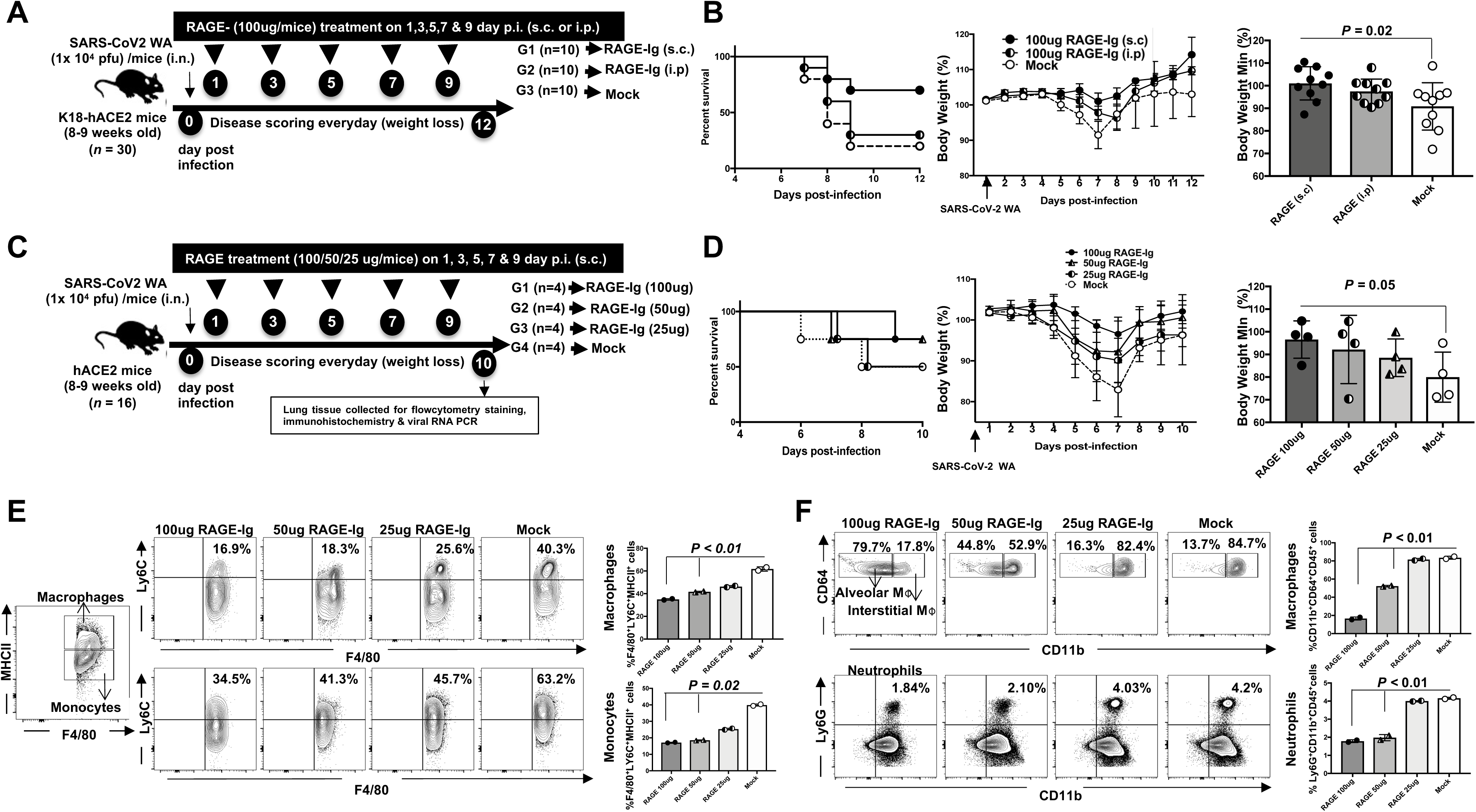
The effect of RAGE-Ig treatment on COVID-like disease and survival in ACE2 mice infected with SARS-CoV-2. (**A**) Experimental scheme to study the effect of route of administration of RAGE-Ig fusion protein on disease outcome and death in ACE2 mice infected with SARS-CoV-2 (Strain Washington, (WA)). Mice are infected with SARS-CoV-2 (WA strain) and left untreated (Mock, *n* = 10,) or treated with RAGE-Ig s.c. (*n* = 10) or i.p. (*n* = 10). (**B**) The graph in the *left panel* shows the percentages of survival of ACE2 mice infected with SARS-CoV-2 and either left untreated (*Mock*) or treated with 100 ug of RAGE-Ig administered subcutaneously (s.c.) or intraperitoneally (i.p.). The graph in the *middle panel* shows the average daily weight change recorded for 12 days calculated in percentages, normalized to the body weight on day 0 of infection. The bar graph in the *right panel* shows the maximal weight change in percentage on day 7 p.i. of ACE2 mice infected with SARS-CoV-2 and either left untreated (*Mock*) or treated with 100 ug of RAGE-Ig administered subcutaneously (s.c.) or intraperitoneally (i.p.). (**C**) Experimental scheme to study the dose-response of RAGE-Ig fusion protein treatment on survival and disease outcome in ACE2 mice infected with SARS-CoV-2. (**D**) The graph in the *left panel* shows the percentages of survival of ACE2 mice infected with SARS-CoV-2 and either left untreated (mock) or treated with different doses of RAGE-Ig administered s.c. The graph in the *middle panel* shows the average daily weight change recorded for 10 days. Bars represent means ± SEM. The bar graph in the *right panel* shows the maximal weight change in percentage on day 7 p.i. of ACE2 mice infected with SARS-CoV-2 and either left untreated (*Mock*) or treated with different doses of RAGE-Ig administered s.c. (**E**) Representative (*left 9 panels*) and average (*right 2 panels*) frequencies of F4/80^+^LY6C^+^MHCII^+^ macrophages (*top panels*) and F4/80^+^LY6C^+^MHCII^-^ monocytes (*bottom panels*) infiltrating the lungs of ACE2 mice infected with SARS-CoV-2 and either left untreated (*Mock*) or treated with 3 different doses of RAGE-Ig. (**F**) Representative (*left 8 panels*) and average (*right 2 panels*) frequencies of macrophage subsets (*top panels*) and neutrophils (*bottom panels*) infiltrating the lungs of ACE2 mice infected with SARS-CoV-2 and either left untreated (*Mock*) or treated with 3 different doses of RAGE-Ig. Bars represent means ± SEM. Data were analyzed by student’s *t*-test comparing RAGE-treated and untreated mice.

### 2. RAGE-Ig treatment protects against COVID-19-like symptoms, virus replication, and mortality in K18-hACE2 mice following infection with various SARS-CoV-2 VOCs

We next determined whether RAGE-Ig treatment would be effective against the wild-type SARS-CoV-2 (USA-WA1/2020) and SARS-CoV-2 VOCs measured by disease outcome and survival in mice. Eight-to-nine-week-old male and female K18-hACE2 transgenic mice were intranasally infected with 1 x 10^4^ pfu of SARS-CoV-2 (USA-WA1/2020) (*n* = 20), B 1.1.7 Alpha (8 x 10^2^ pfu) (*n* = 10), B.1.351 Beta (6 x 10^3^ pfu) (*n* = 10), and P. 1 Gamma (2 x 10^2^ pfu) (*n* = 10) variants. Subsequently, mice were treated subcutaneously (s.c.) with 100 μg RAGE-Ig/mouse or mock-treated with a buffer on alternate days (1, 3, 5, 7, and 9 days p.i.) (**Fig. 2A**). RAGE-Ig treatment (100 μg) was found to significantly protective against weight loss, mortality, and viral load in USA-WA1/2020, B.1.351 Beta, and P. 1 Gamma-infected mice (**Fig. 2B**). H & E staining of lung sections demonstrated a reduction in lung pathogenicity in the RAGE-Ig-treated mice compared to mock (vehicle-treated) mice at day 14 p.i. (**Fig. 2C**). The most surprising result was a decrease in viral RNA in the lungs of RAGE-Ig-treated mice compared to mock-treated mice (**Fig. 2B**), suggesting a possible anti-viral property of the RAGE-Ig compound.

**Figure 2.**
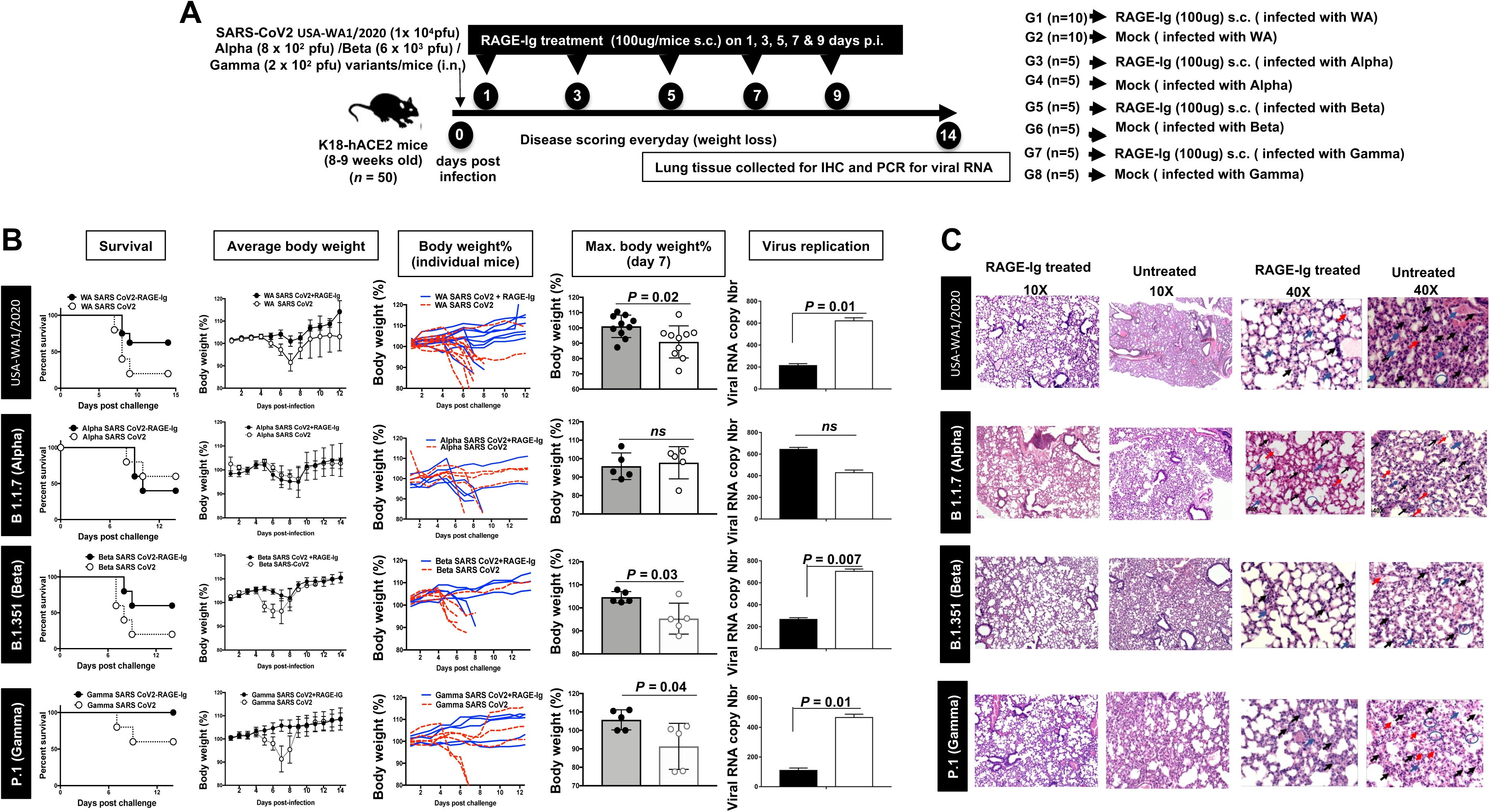
The effect of RAGE-Ig treatment on disease outcome and survival during SARS-CoV-2 VoCs infection in mice. (**A**) Experimental scheme to understand the effect of RAGE-Ig treatment on survival, disease outcome, and virus replication upon infection with the SARS-CoV-2 variants. (**B**) Mice that are infected with SARS-CoV-2 and then treated with RAGE-Ig are represented as a closed black circle and mock-infected as an open white circle. The left panel shows the survival graph for each variant of SARS-CoV2. The line graph (*second to panel*) shows the average weight change of 10 mice at each day p.i. normalized to the body weight on the day of infection for each variant of concern. The line graph (*third to left panel*) shows weight change for individual mice at each day p.i. normalized to the body weight on the day of infection for each variant. Bars represent mean ± SEM of weight change at day 7 p.i. in percentage, normalized to the body weight on the day of infection. The bar graph (right panel) shows virus replication in the lungs of RAGE-Ig-treated and untreated mice. The graph indicated fold change viral RNA copy numbers for each variant of concern. On day 14 p.i., mice are euthanized, and lungs are harvested and used for viral RNA quantification by qRT-PCR. (**C**) The H&E images show representative inflammatory cells infiltrating the lungs of mice infected with SARS-CoV-2 variants and treated with RAGE-Ig. At day 12 p.i., hamsters were euthanized, and lungs are analyzed by immunohistochemistry for pathology. The H&E images are shown at a magnification of 10X (left two panels) and 40X (right two panels). Bars represent mean ± SEM. Data analyzed by student’s *t*-tests compares RAGE-treated and untreated mice.

### 3. RAGE-Ig treatment protects golden Syrian hamsters from COVID-19 symptoms and virus replication following infection with SARS-CoV-2 WA and Delta variants

Although the protective effect of RAGE-Ig treatment against wild-type WA or VOCs infection was confirmed using the mouse model, we also evaluated RAGE-Ig in the hamster model, since hamsters are outbred and recapitulate the SARS-CoV-2 pathogenicity with a higher degree of similarity to humans. Seven to eight weeks old male Syrian golden hamsters were intranasally infected with 1 x 10^5^ pfu of SARS-CoV-2 (USA-WA1/2020) (*n* = 10) or B.1.617.2 Delta (6.9 x 10^4^ pfu) (*n* = 10)/hamster in 100 μl and followed by treatment with 1.5 mg RAGE-Ig/hamster (*n* = 3) or 3 mg RAGE-Ig/hamster (*n* = 3) by either the subcutaneous (s.c.) route or mock-treated with buffer (*n* = 4) on alternate days (1, 3, 5, 7 and 9 days p.i.) **Fig. 3A**. Hamsters infected with SARS-CoV-2 WA or Delta viruses and treated with RAGE-Ig experienced significantly less weight loss than the mock-treated group (**Fig. 3B**). We found reduced pathogenicity and viral load in RAGE-Ig-treated hamsters, similar to the results observed in mice. There was a decrease in viral RNA in the lungs of RAGE-Ig-treated hamsters as compared to mock-treated hamsters (**Fig. 3B**). Thus, our results in the hamster model confirmed the protective anti-viral effect of the drug against SARS-CoV-2 infection across different models of infection.

**Figure 3.**
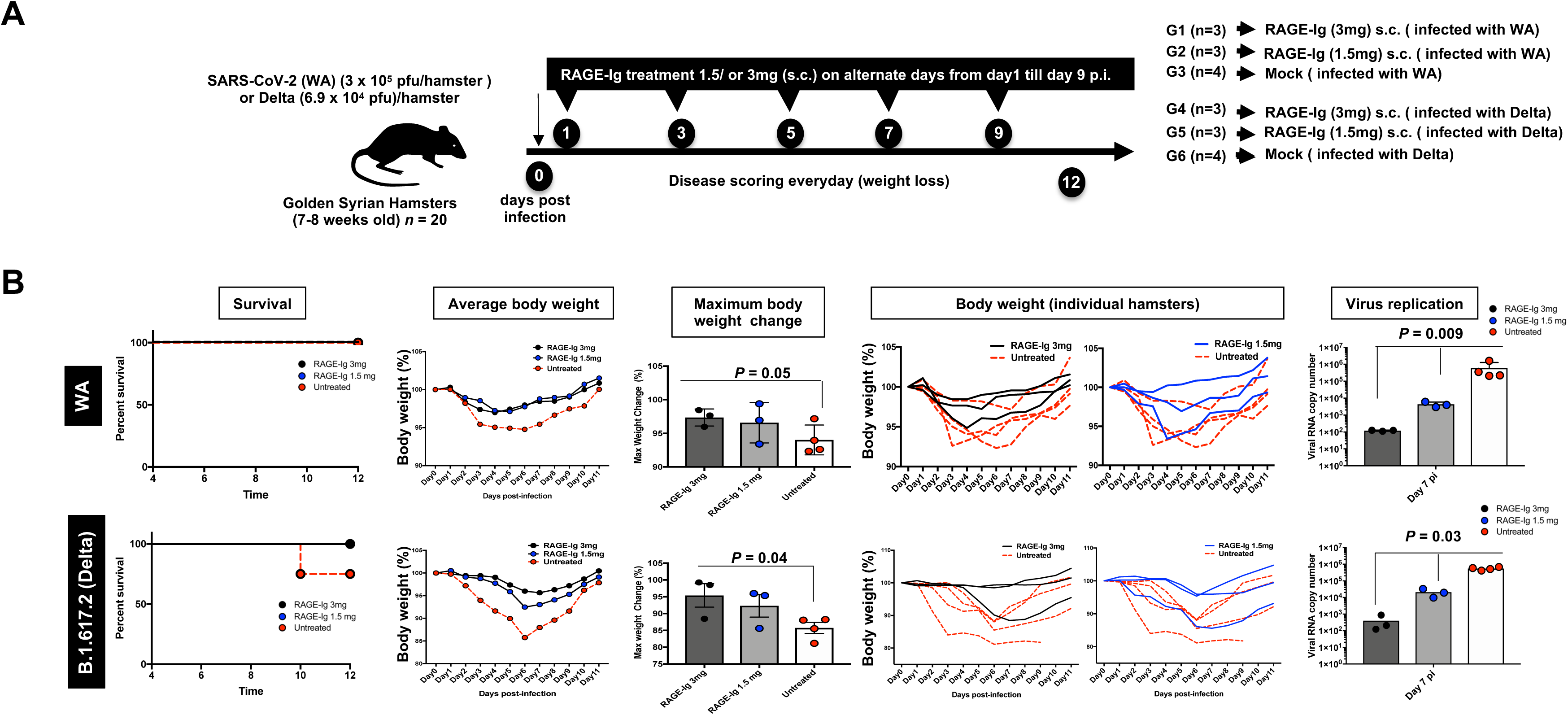
Effect of RAGE-Ig treatment on disease outcome and survival in the golden Syrian hamster infected with SARS-CoV-2 variants. (**A**) Experimental scheme to study the effect of RAGE-Ig treatment on survival, disease outcome, and virus replication in hamsters infected with SARS-CoV-2 WA or delta variants. (**B**) Hamsters infected with SARS-CoV-2 and treated with RAGE-Ig are represented as a closed black circle (3 mg) or blue circle (1.5 mg). Mock-treated hamsters are represented as an open circle. The graph in the left panel shows the survival of treated and untreated hamsters for the WA variant(*top*) and Delta variant (*bottom*). The line graph (*second to left panel*) shows weight change for 14 days p.i. normalized to the body weight on the day of infection for each variant. The bar graph shows the percentage of maximum weight loss in RAGE-Ig treated and untreated hamsters on day 8 post-infection. Individual line graph (fourth and fifth panels from the left) showing weight change in individual hamsters for 11 days p.i. normalized to the body weight on the day of infection with the WA variant (*top panel*) and Delta variant (*bottom panel*). The bar graph (right panel) shows virus replication in the lungs of RAGE-Ig-treated and untreated hamsters. The graph a fold change in viral RNA copy numbers for each variant as indicated. At day 12 p.i., hamsters were euthanized, and lungs were used for viral RNA quantification by qRT-PCR. Bars represent mean ± SEM. Data analyzed by student’s *t*-test compares RAGE-treated and untreated hamsters.

### 4. The anti-viral effect of RAGE-Ig in SARS-CoV-2 infected human lung epithelial cells is mediated by type I and type III interferons

Our results from the mouse and hamster models revealed an anti-viral effect of RAGE-Ig, which is host-induced. Interferons are the major anti-viral players during SARS-CoV-2 infection; hence we studied the effect of RAGE-Ig treatment on Type I, II & III interferon secretion in SARS-CoV-2 infected human lung epithelial cells (BEAS-2B) and its anti-viral effect in infected lungs (**Supplemental Fig. S2)**. BEAS-2B cells were infected with SARS-CoV-2 USA-WA1/2020 and treated with various concentrations of RAGE-Ig (30,100, 250, and 500 ng/ml). Supernatants and cell lysates were collected at various time points (24, 48, and 72 hours) for ELISA and qRT-PCR (**Fig. 4A**). There was a statistically significant decrease in viral RNA copy number at 24 (left panel) and 48 hours (right panel) in the lysates of BEAS-2B cells infected with SARS-CoV-2 following RAGE-Ig treatment as quantified by qRT-PCR (**Fig. 4B**), thus confirming the anti-viral effect of RAGE-Ig *in vitro*. Simultaneously, there was an increased expression of the interferon Type I and III genes at 24 and 48 hours in BEAS-2B cell lysates infected with SARS-CoV-2 following treatment with RAGE-Ig (**Fig. 4C**). In addition, we detected an increase in interferon beta, gamma, and lambda3 cytokine release at 48 hours p.i. in the lysates of infected BEAS-2B cells, following treatment with RAGE-Ig. From this, it may be inferred that induced anti-viral interferons, particularly Type I and III, can curtail SARS-CoV-2 multiplication in human lung epithelial cells.

**Figure 4.**
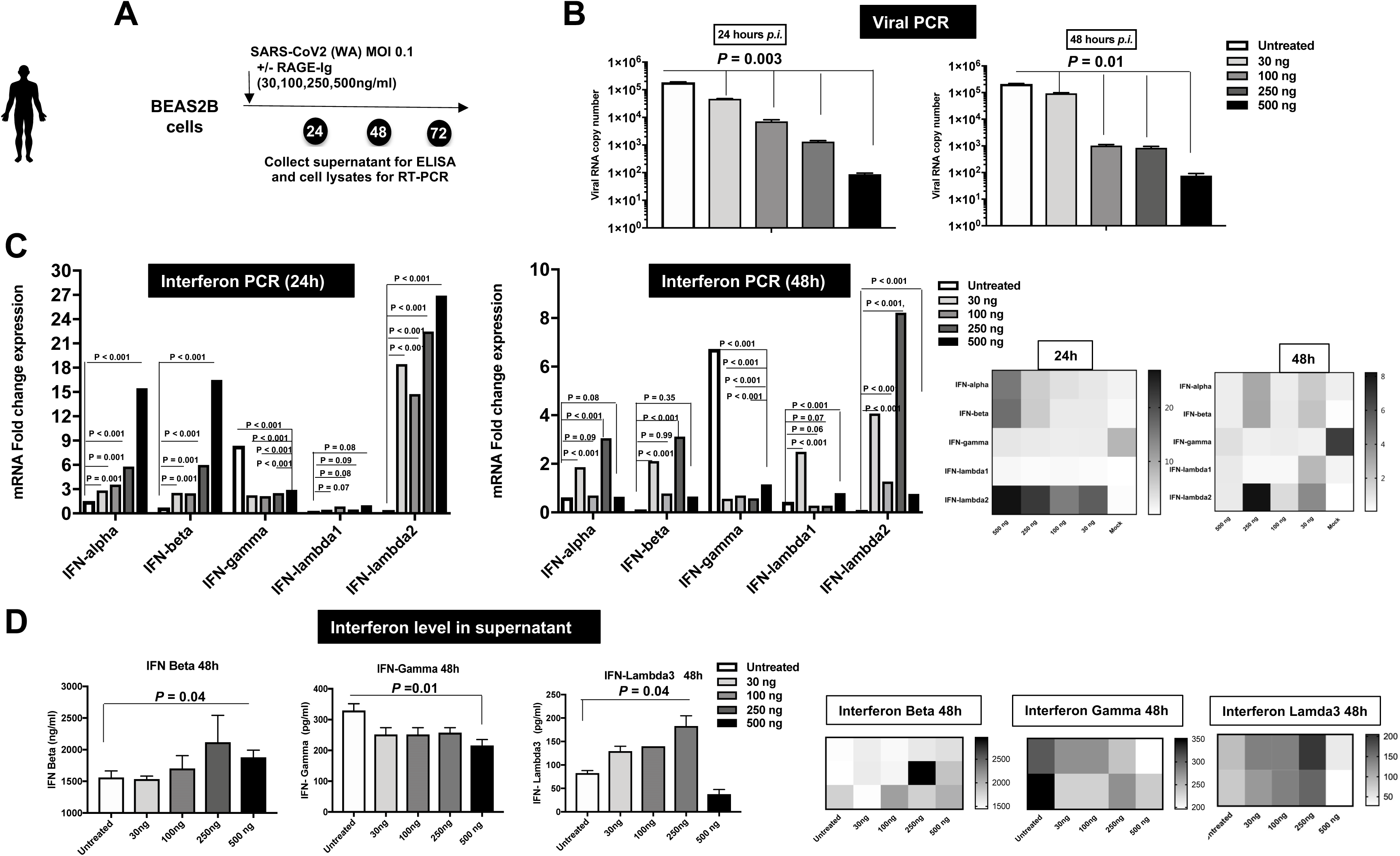
Effect of RAGE-Ig treatment on interferon type I, II & III interferon production *in vitro* by SARS-CoV-2 infected human lung epithelial cells. (**A**) Experimental scheme to study the effect of RAGE-Ig treatment on production of type I, II & III interferon in SARS-CoV-2 infected human lung epithelial cells. The BEAS-2B human lung epithelial cells are infected with SARS-CoV-2 WA and then treated with various doses of RAGE-Ig (30, 100, 250 & 500 ng/ml). Supernatant and cell lysates were collected at various time points (24, 48, and 72 hours). (**B**) The graph shows viral RNA copy numbers at 24 hours (*left panel*) and 48 hours (*right panel*) in the lysates of BEAS-2B cells infected with SARS-CoV-2 and treated with RAGE-Ig. (**C**) The graph shows mRNA expression of interferon type I, II, and III genes at 24 hours (*left panel*) and 48 hours (*middle panel*) in the lysates of BEAS-2B cells infected with SARS-CoV-2 and treated with RAGE-Ig. The corresponding heat map is shown in the right panel. (**D**) Graphs show the amount of interferon beta, gamma, and lambda 3 released 48 hours in the lysates of BEAS-2B cells infected with SARS-CoV-2 and treated with RAGE-Ig (left panels). Bars represent mean ± SEM. The corresponding heat map for interferon beta, gamma, and lambda3 release is shown in the right panels. Data analyzed by student’s t-test compares RAGE-treated and untreated cells.

### 5. The anti-inflammatory effects of RAGE-Ig in SARS-CoV-2 infected human lung epithelial cells infected with SARS-CoV-2

The potential therapeutic effect of RAGE-Ig protein against SARS-CoV-2 infection in mice is partly due to a decrease in macrophage-related inflammatory cell infiltration. Although the anti-inflammatory effect of RAGE-Ig is well-known, we wanted to re-demonstrate this effect against SARS-CoV-2 *in vitro*. To confirm the anti-inflammatory effect of RAGE-Ig treatment, BEAS-2B cells were infected with SARS-CoV-2 USA-WA1/2020 followed by treatment with various concentrations of RAGE-Ig (30, 100, 250, and 500 ng/ml). Supernatants were collected at multiple time points (24, 48, and 72 hours) for cytokine estimation. Kinetics of the IL-6 and IL-8 cytokines released in the supernatants of BEAS-2B cells infected with SARS-CoV-2 showed a maximal release at 48 hours p.i. (**Fig. 5A**). Additionally, we observed a significant dose-dependent decrease in the release of IL-6 and IL-8 cytokines at 48 hours in the supernatants of BEAS-2B cells infected with SARS-CoV-2 following treatment with RAGE-Ig (**Fig. 5B**). Similarly, there was a dose-dependent decrease in the levels of IL-6 and IL-8 cytokines at 48 hours in the supernatants of human PBMCs infected with SARS-CoV-2 (**Fig. 5C**). RAGE-Ig treatment *ex vivo* conferred anti-inflammatory effects both on the inflammatory cells from COVID-19 patients PBMCs and on human lung epithelial cells infected with SARS-CoV-2 *in vitro*.

**Figure 5.**
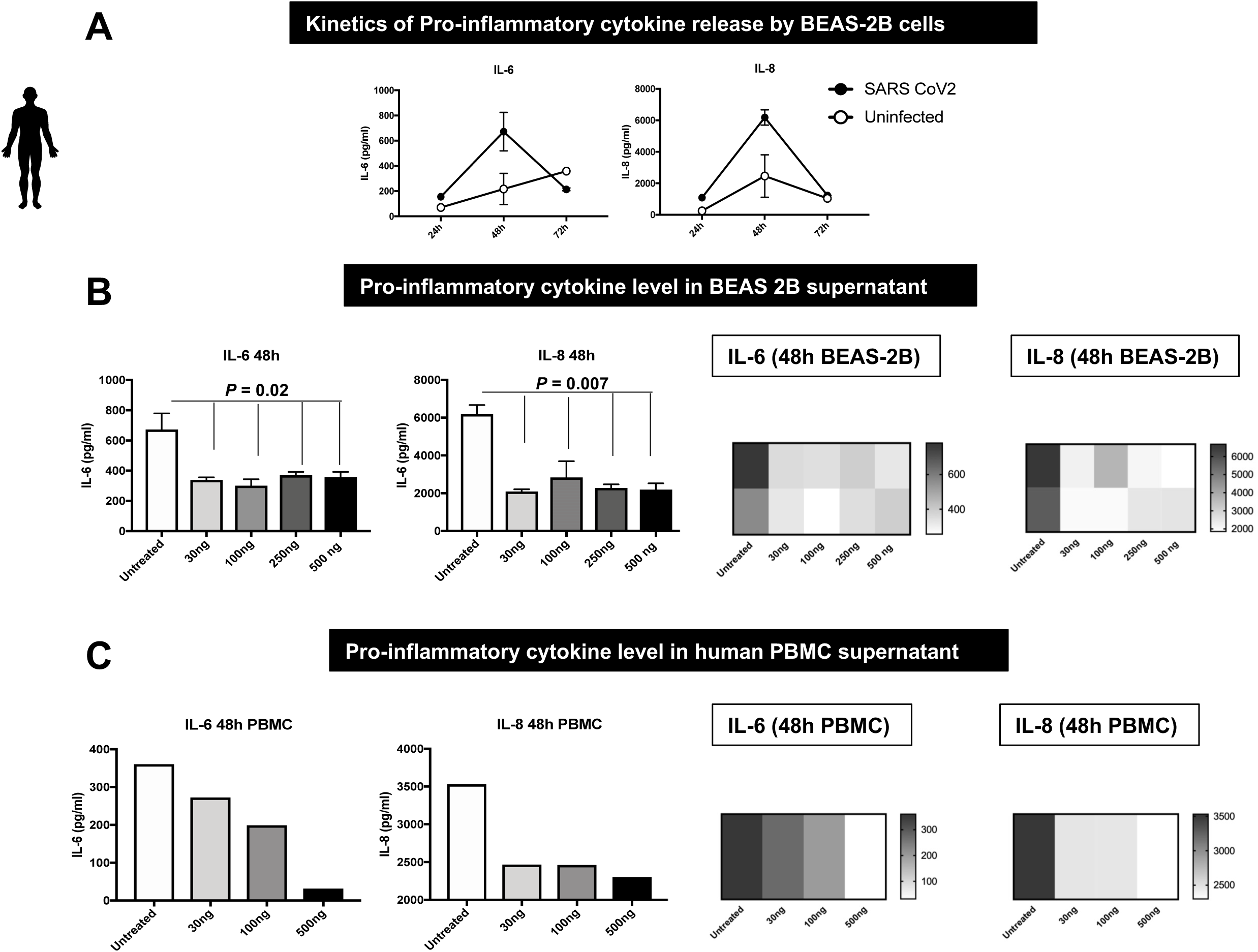
Effect of RAGE-Ig treatment on production of pro-inflammatory cytokines *in vitro* by SARS-CoV-2 infected human lung epithelial cells. BEAS-2B cells are infected with SARS-CoV-2 (WA strain) and then treated with various concentrations of RAGE-Ig (30, 100, 250 & 500 ng/ml). Supernatants were collected at various time points (24, 48, and 72 hours) and the level of cytokine was determined by Luminex. (**A**) Graphs show the kinetics of IL-6 (*left*) and IL-8 (*right*) cytokines released in the supernatants of BEAS-2B cells infected with SARS-CoV-2. (**B**) Graphs show the effect of RAGE-Ig treatment on the release of IL-6 and IL-8 cytokines at 48 hours in the supernatants of BEAS-2B cells infected with SARS-CoV-2 (*left 2 panels).* The corresponding heat map for cytokine release is shown in the *right 2 panels.* (**C**) Graphs show the effect of RAGE-Ig treatment on the release of IL-6 and IL-8 cytokines at 48 hours in the supernatants of human PBMC infected with SARS-CoV-2 (*left 2 panels)*. The corresponding heat map for cytokine release is shown in the *right 2 panels.* Bars represent means ± SEM. Data analyzed by student *t*-test compares RAGE-treated and untreated cells.

### 6. RAGE-Ig treatment affects the inflammatory profile of monocytes from COVID-19 patients

The interaction between RAGE and its ligands may ultimately activate the pro-inflammatory gene. PBMCs from asymptomatic (ASYMP) and symptomatic (SYMP) COVID-19 patients were stained for the expression of RAGE ligand EN-RAGE on CD14^+^ monocytes *ex vivo* (**Fig. 6A**). To examine if RAGE-Ig treatment can modulate the expression of inflammatory receptors, PBMCs from SYMP COVID-19 patients were treated with various concentrations of RAGE-Ig (10 μg, 1 μg, 100 ng, and 10 ng/ml) and analyzed by flow cytometry. RAGE-Ig treatment significantly decreased the expression of CD64 on monocytes of COVID-19 patients in a dose-dependent manner (**Fig. 6B**). CD64 is a high-affinity IgG receptor (FcγRI) that contributes to COVID-19-associated inflammation severity. Thus, RAGE-Ig treatment can potentially alleviate immune complex-mediated airway inflammation. Moreover, CD64^-^ monocytes have been implicated in anti-viral immunity. To further understand whether RAGE-Ig treatment can modulate macrophage differentiation, human CD14^+^ monocytes were isolated from PBMCs and differentiated into uncommitted macrophages (M0) by macrophage colony-stimulating factor (M-CSF). We found that the RAGE-Ig protein treatment caused dose-dependent morphological changes in M0 macrophages during the differentiation process as shown in the figure (**Fig. 6C**).

**Figure 6.**
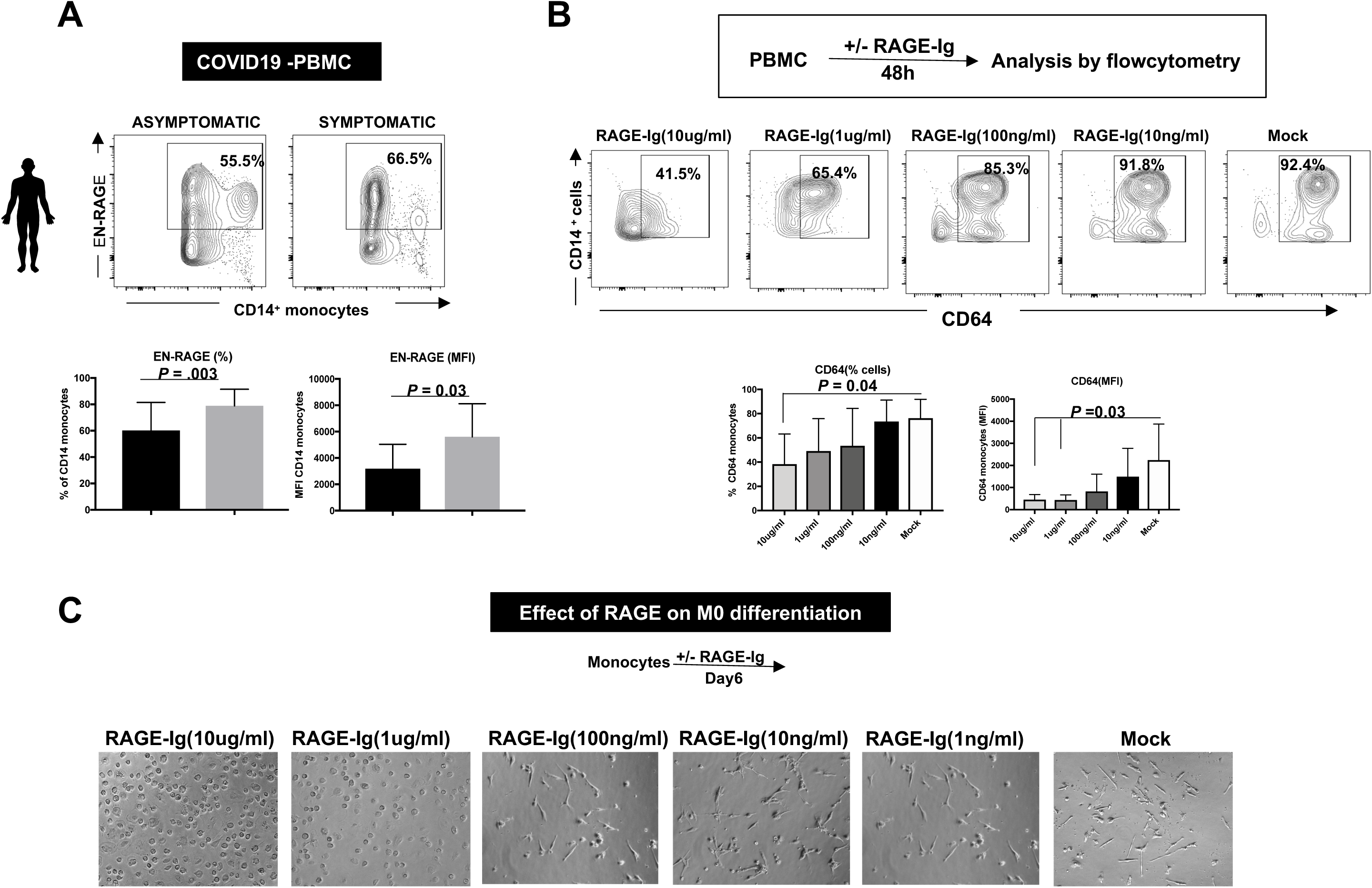
Effect of RAGE-Ig treatment on expression of inflammatory receptors on monocytes from COVID-19 patients. (**A**) The expression of RAGE ligand EN-RAGE on CD14^+^ monocytes *ex-vivo* by PBMCs from symptomatic (SYMP) and asymptomatic (ASYMP) COVID-19 patients detected by FACS. The representative (top panels) and average (bottom panels) data of EN-RAGE expression on CD14^+^ monocytes in ASYMP and SYMP COVID-19 patients are shown. (**B**) PBMCs from SYMP COVID-19 patients were treated with various concentrations of RAGE-Ig (10 ug, 1ug, 100ng, 10ng/ml), and the frequencies of CD14^+^CD64^+^ monocytes are analyzed by flow cytometry. The representative (top panels) and average (bottom panels) frequencies of CD14^+^CD64^+^ monocytes upon RAGE-Ig treatment are shown. The bottom right panel shows the effect of RAGE-Ig treatment on the level of expression of CD64 in monocytes from COVID-19 patients. (**C**) Representative microscopic images showing dose-dependent morphological changes in M0 macrophages during monocyte differentiation into a macrophage after RAGE-Ig protein treatment. Human CD14^+^ monocytes were isolated from PBMC and further differentiated into uncommitted macrophages (M0) by macrophage colony-stimulating factor (M-CSF) and then treated with RAGE-Ig protein. Bars represent means ± SEM. Data were analyzed by student’s t-test comparing RAGE-treated and untreated monocytes.

## DISCUSSION

Anti-inflammatory drug interventions can be critical in preventing complications of acute COVID-19 and shortening disease recovery time in long-COVID-19 patients. Both clinical and epidemiological evidence of COVID-19 associates age and different comorbidities with the risk of SARS-CoV-2 infection leading to severe lung involvement and disease. The association of the RAGE pathway with aging, and COVID-19-associated comorbidities like diabetes, hypertension, CAD, CVD, renal diseases, and others, suggest that intervention of the RAGE pathway may potentially be a promising therapeutic target for treating patients with pre-existing conditions and severe COVID-19 symptoms (*19–29*). Interestingly, a recent study has reported high expression of RAGE and HMGB1 in the lung tissue of decedents with COVID-19 and diabetes. The use of RAGE inhibitors thus may be a novel therapeutic target in the prevention, or slowing of the progression of SARS-CoV-2 infections that currently lack effective therapy. In this study, we explored the potential use of Galactica’s RAGE-Ig protein as a therapeutic intervention to attenuate inflammatory reactions in possible approaches for treatment against SARS-CoV-2 infection with VOCs. For this, we first tested the drug’s therapeutic effect against the wild-type WA variant of SARS-CoV-2 infection in K18-hACE2 transgenic mice. We observed a statistically significant dose-dependent protection on survival and weight loss of K18-hACE2 mice infected with the WA strain of SARS-CoV-2. Importantly, we found that 100 μg of RAGE-Ig administered s.c. was the most effective dose and route of administration that protected against the symptoms of COVID-19 in mice. Subsequent experiments demonstrated that RAGE-Ig-treated mice had significantly decreased weight loss and mortality compared to mock-treated K18-hACE2 mice infected with Alpha, Beta, or Gamma variants of SARS-CoV-2. A recent study has reported the effect of RAGE antagonist treatment in a mouse model to be effective against COVID-19 (*36*). To our knowledge, this is the first study to report the effect of RAGE protein on SARS-CoV-2 VOCs.

For pre-clinical testing, we confirmed the potential therapeutic effect of this drug at a multi-species level. We tested the drug’s effect in Syrian golden hamsters infected with WA or Delta VOCs of SARS-CoV-2. We demonstrated that the hamsters were protected from the symptoms of SARS-CoV-2 VOCs infection when treated with 1.5 or 3 mg RAGE-Ig/hamster/dose. This pre-clinical data demonstrated that RAGE-Ig treatment can be clinically useful in preventing complications of COVID-19 acute infection and shortening disease recovery time. Hospitalized COVID-19 patients requiring ventilation or expected to do so imminently will be the most important clinical criteria indicating treatment with RAGE-Ig. Survival among these patients will be the primary endpoint tested in this compound’s COVID-19 clinical trial.

The anti-inflammatory effect of the RAGE-Ig is well-established in the context of other chronic inflammatory diseases. Therefore, we intended to further understand it specifically in the context of COVID-19 infection. Due to an enhanced level of RAGE ligands in diabetes or other chronic disorders, this receptor has a causative effect in a range of inflammatory diseases. Given its inflammatory function and the ability to detect multiple ligands through a common structural motif, RAGE is often referred to as a pattern recognition receptor. EN-RAGE is a ligand for the receptor for advanced glycation end products (RAGE). Prior studies have demonstrated that the expression of RAGE ligand S100A12 (also known as EN-RAGE), a biomarker of pulmonary injury, increases in peripheral monocytes of COVID-19 patients. Increased EN-RAGE may promote pulmonary damage in these patients by activation of RAGE receptors on the surface of alveolar epithelial cells. We next examined the role played by RAGE and its ligands in the disease severity of COVID-19 patients, by analyzing monocytes from PBMCs of COVID-19 patients for expression of ligands and receptors of RAGE and correlated with disease severity. Instead of previous reports, we observed an increased expression of EN-RAGE in symptomatic cases of COVID-19 compared to asymptomatic COVID-19 patients. In our study, we detected decreased expression of CD64(FcgR1) on monocytes upon treatment with RAGE-Ig. CD64 is a high-affinity IgG receptor (FcγRI) that contributes to inflammation severity in multiple disease models. Thus, RAGE-Ig treatment can potentially alleviate immune complex-mediated airway inflammation. Moreover, CD64^-^ monocytes have been implicated in anti-viral immunity. Circulating monocytes differentiate upon entering tissues and are activated by cytokines or growth factors present in the lung during active infection.

Activation of uncommitted macrophages (M0) classically polarizes them into either pro-inflammatory M1 macrophages or anti-inflammatory M2 macrophages. Interestingly, our results showed that RAGE-Ig treatment caused a dose-dependent morphological change in M0 macrophages *in vitro,* thus tempting speculation that the compound can affect macrophage polarization. Upon infection, monocytes migrate into the tissues, where they become infected resident macrophages, allowing viruses to spread within tissues and organs, such as the lungs. SARS-CoV-2-infected lung-resident pro-inflammatory M1 macrophage produces large amounts of pro-inflammatory cytokines and chemokines, known as cytokine storm, leading to an excessive, uncontrolled local tissue inflammatory response with tissue damage. All this could result in acute respiratory illness, known as severe acute respiratory syndrome (SARS), characterized by fever, productive cough, shortness of breath/dyspnea, and pneumonia-like symptoms. Both local tissue inflammation and the cytokine storm play an immune-pathological role in developing COVID-19-related complications, such as ARDS, the main cause of death in COVID-19 patients. A molecular treatment that would reduce the development of pro-inflammatory M1 macrophages or directly inhibit the production of pro-inflammatory cytokines and chemokines is urgently needed to reduce ARDS. Interestingly, we also observed a dose-dependent decreased infiltration of neutrophils (CD11b/Ly6G) and inflammatory macrophages (F4/80/Ly6C) following treatment with RAGE-Ig in SARS-CoV-2 infected mouse lung samples. In addition, we also found decreased Interstitial/exudative macrophages in SARS-CoV-2 infected mouse lungs treated with RAGE-Ig.

The current protocols for the treatment of COVID-19 are largely based on the use of anti-inflammatory compounds like dexamethasone, baricitinib and/or monoclonal antibodies, Molnupiravir and Nirmatrelvir/Ritonavir, etc. There is a critical need for other medicinal agents, especially those with dual antiviral and anti-inflammatory (AAI) activity, that would be readily available for the early treatment of mild to moderate COVID-19 in high-risk patients. Interestingly, we noticed that RAGE-Ig treatment not only protected from symptoms of SARS-CoV-2 infection but also decreased the viral titers of SARS-CoV-2 in the lungs of treated animals. This will greatly improve the survival outcomes of ventilated COVID-19 patients by shortening their ICU stays and reducing their viral load. Although the underlying causes of long COVID-19, which may affect at least ten percent of all those infected with COVID-19, are not yet well understood, clinicians believe that long COVID-19 symptoms are caused by residual inflammation and/or viral load. If this hypothesis is correct, the RAGE-Ig protein may also be an effective therapy for long COVID-19 with its unique dual anti-inflammatory and antiviral activity.

Due to its rare dual properties of antiviral and anti-inflammatory activity (AAI), the RAGE-Ig protein, with its broad preclinical efficacy, is a promising drug candidate for COVID-19 treatment. To confirm whether the RAGE-Ig treatment induced any anti-viral effect, we conducted a series of in vitro studies of the drug using a human lung epithelial cell line (BEAS-2B). We demonstrated the role of the Interferon pathway in the anti-viral mechanisms induced during RAGE-Ig treatment of lung epithelial cells. RAGE-Ig treatment of SARS-CoV-2 infected epithelial cells caused a dose-dependent decrease in viral RNA copy number. This, in turn, was found to correspond to an increase in Type I and Type III interferons, as demonstrated by both qRT-PCR and ELISA. our findings suggest that RAGE-Ig treatment increased Type I & III interferon but not Type II interferon (IFN-Gamma) response in human lung epithelial cells infected with SARS-CoV-2.

It is established that Type I interferon signaling leads to anti-viral ISG expression (via IRF3 and IRF7) whereas Type II interferon signaling leads to pro-inflammatory gene expression (via NFkB signaling). An unbalanced immune response, characterized by a weak production of Type I interferons (IFN-Is) and an exacerbated release of pro-inflammatory cytokines, contributes to the severe forms of the disease. Thus, RAGE-Ig treatment during SARS-CoV-2 infection may suppress inflammatory cytokine responses (such as IL-6 and IL-8) and enhance Type I and III interferons (IFN-Beta and IFN-lambda) response resulting in less severe disease. Our results indicate a dual antiviral and anti-inflammatory (AAI) activity and significant therapeutic potential for Galactica’s RAGE-Ig protein against COVID-19 caused by emerging VOCs. The therapeutic effect of RAGE-Ig on SARS-CoV-2 infection is mediated through anti-inflammatory as well as a host-directed anti-viral effect by Type I and Type III interferons.

## ACKNOWLEDGEMENTS

This work is supported by a Grant from Galactica Pharmaceuticals, Inc., Villanova, PA 19085; by the Fast-Grant PR12501 from Emergent Ventures, and by Public Health Service Research grants AI158060, AI150091, AI143348, AI147499, AI143326, AI138764, AI124911, and AI110902 from the National Institutes of Allergy and Infectious Diseases (NIAID) to LBM and by R43AI174383-01 to TechImmune, LLC.

## REFERENCES

1. A. Lim, A. Radujkovic, M. A. Weigand, U. Merle, Soluble receptor for advanced glycation end products (sRAGE) as a biomarker of COVID-19 disease severity and indicator of the need for mechanical ventilation, ARDS and mortality. Ann Intensive Care 11, 50 (2021).

2. D. Yalcin Kehribar et al., The receptor for advanced glycation end product (RAGE) pathway in COVID-19. Biomarkers 26, 114–118 (2021).

3. S. Chiappalupi et al., Targeting RAGE to prevent SARS-CoV-2-mediated multiple organ failure: Hypotheses and perspectives. Life Sci 272, 119251 (2021).

4. M. Jabaudon et al., Receptor for advanced glycation end-products and ARDS prediction: a multicentre observational study. Sci Rep 8, 2603 (2018).

5. M. Kerkeni, J. Gharbi, RAGE receptor: May be a potential inflammatory mediator for SARS-COV-2 infection? Med Hypotheses 144, 109950 (2020).

6. M. Salehi, S. Amiri, D. Ilghari, L. F. A. Hasham, H. Piri, The Remarkable Roles of the Receptor for Advanced Glycation End Products (RAGE) and Its Soluble Isoforms in COVID-19: The Importance of RAGE Pathway in the Lung Injuries. Indian J Clin Biochem, 1–13 (2022).

7. A. Bierhaus et al., Understanding RAGE, the receptor for advanced glycation end products. J Mol Med (Berl) 83, 876–886 (2005).

8. E. A. Oczypok, T. N. Perkins, T. D. Oury, All the “RAGE” in lung disease: The receptor for advanced glycation endproducts (RAGE) is a major mediator of pulmonary inflammatory responses. Paediatr Respir Rev 23, 40–49 (2017).

9. C. S. Calfee et al., Plasma receptor for advanced glycation end products and clinical outcomes in acute lung injury. Thorax 63, 1083–1089 (2008).

10. T. Nakamura et al., Increased levels of soluble receptor for advanced glycation end products (sRAGE) and high mobility group box 1 (HMGB1) are associated with death in patients with acute respiratory distress syndrome. Clin Biochem 44, 601–604 (2011).

11. M. Jabaudon et al., Soluble Receptor for Advanced Glycation End-Products Predicts Impaired Alveolar Fluid Clearance in Acute Respiratory Distress Syndrome. Am J Respir Crit Care Med 192, 191–199 (2015).

12. R. Blondonnet et al., RAGE inhibition reduces acute lung injury in mice. Sci Rep 7, 7208 (2017).

13. Y. S. Sim, D. G. Kim, T. R. Shin, The diagnostic utility and tendency of the soluble receptor for advanced glycation end products (sRAGE) in exudative pleural effusion. J Thorac Dis 8, 1731–1737 (2016).

14. A. Sharma, S. Kaur, M. Sarkar, B. C. Sarin, H. Changotra, The AGE-RAGE Axis and RAGE Genetics in Chronic Obstructive Pulmonary Disease. Clin Rev Allergy Immunol 60, 244–258 (2021).

15. C. Machahua et al., Serum AGE/RAGEs as potential biomarker in idiopathic pulmonary fibrosis. Respir Res 19, 215 (2018).

16. J. Tian et al., Toll-like receptor 9-dependent activation by DNA-containing immune complexes is mediated by HMGB1 and RAGE. Nat Immunol 8, 487–496 (2007).

17. M. A. Hofmann et al., RAGE and arthritis: the G82S polymorphism amplifies the inflammatory response. Genes Immun 3, 123–135 (2002).

18. U. Ekong et al., Blockade of the receptor for advanced glycation end products attenuates acetaminophen-induced hepatotoxicity in mice. J Gastroenterol Hepatol 21, 682–688 (2006).

19. K. Kuba, Y. Imai, J. M. Penninger, Angiotensin-converting enzyme 2 in lung diseases. Curr Opin Pharmacol 6, 271–276 (2006).

20. J. M. Forbes et al., Modulation of soluble receptor for advanced glycation end products by angiotensin-converting enzyme-1 inhibition in diabetic nephropathy. J Am Soc Nephrol 16, 2363–2372 (2005).

21. E. Dozio et al., Soluble Receptor for Advanced Glycation End Products and Its Forms in COVID-19 Patients with and without Diabetes Mellitus: A Pilot Study on Their Role as Disease Biomarkers. J Clin Med 9, (2020).

22. E. M. De Francesco, V. Vella, A. Belfiore, COVID-19 and Diabetes: The Importance of Controlling RAGE. Front Endocrinol (Lausanne) 11, 526 (2020).

23. A. Rojas, I. Gonzalez, M. A. Morales, SARS-CoV-2-mediated inflammatory response in lungs: should we look at RAGE? Inflamm Res 69, 641–643 (2020).

24. R. S. Stilhano et al., SARS-CoV-2 and the possible connection to ERs, ACE2, and RAGE: Focus on susceptibility factors. FASEB J 34, 14103-14119 (2020).

25. S. J. Loomis et al., Cross-sectional Analysis of AGE-CML, sRAGE, and esRAGE with Diabetes and Cardiometabolic Risk Factors in a Community-Based Cohort. Clin Chem 63, 980–989 (2017).

26. F. M. Gutierrez-Mariscal et al., Reduction in Circulating Advanced Glycation End Products by Mediterranean Diet Is Associated with Increased Likelihood of Type 2 Diabetes Remission in Patients with Coronary Heart Disease: From the Cordioprev Study. Mol Nutr Food Res 65, e1901290 (2021).

27. K. Sebekova, Z. Krivosikova, M. Gajdos, Total plasma Nepsilon-(carboxymethyl)lysine and sRAGE levels are inversely associated with a number of metabolic syndrome risk factors in non-diabetic young-to-middle-aged medication-free subjects. Clin Chem Lab Med 52, 139–149 (2014).

28. M. A. Farhangi, P. Dehghan, N. Namazi, Prebiotic supplementation modulates advanced glycation end-products (AGEs), soluble receptor for AGEs (sRAGE), and cardiometabolic risk factors through improving metabolic endotoxemia: a randomized-controlled clinical trial. Eur J Nutr 59, 3009–3021 (2020).

29. H. Koyama et al., Comparison of effects of pioglitazone and glimepiride on plasma soluble RAGE and RAGE expression in peripheral mononuclear cells in type 2 diabetes: randomized controlled trial (PioRAGE). Atherosclerosis 234, 329–334 (2014).

30. L. S. Y. Wong et al., Age-Related Differences in Immunological Responses to SARS-CoV-2. J Allergy Clin Immunol Pract 8, 3251–3258 (2020).

31. H. J. Kim, M. S. Jeong, S. B. Jang, Molecular Characteristics of RAGE and Advances in Small-Molecule Inhibitors. Int J Mol Sci 22, (2021).

32. A. Bierhaus, P. P. Nawroth, Multiple levels of regulation determine the role of the receptor for AGE (RAGE) as common soil in inflammation, immune responses and diabetes mellitus and its complications. Diabetologia 52, 2251–2263 (2009).

33. M. Neeper et al., Cloning and expression of a cell surface receptor for advanced glycosylation end products of proteins. J Biol Chem 267, 14998–15004 (1992).

34. H. J. Huttunen, C. Fages, H. Rauvala, Receptor for advanced glycation end products (RAGE)-mediated neurite outgrowth and activation of NF-kappaB require the cytoplasmic domain of the receptor but different downstream signaling pathways. J Biol Chem 274, 19919–19924 (1999).

35. C. H. Yeh et al., Requirement for p38 and p44/p42 mitogen-activated protein kinases in RAGE-mediated nuclear factor-kappaB transcriptional activation and cytokine secretion. Diabetes 50, 1495–1504 (2001).

36. F. Jessop, et al., Impairing RAGE signaling promotes survival and limits disease pathogenesis following SARS-CoV-2 infection in mice. JCI Insight 7, (2022).

37. J. B. Case, A. L. Bailey, A. S. Kim, R. E. Chen, M. S. Diamond, Growth, detection, quantification, and inactivation of SARS-CoV-2. Virology 548, 39–48 (2020).

