## Supplementary material for "Anti-Viral and Anti-Inflammatory Therapeutic Effect of RAGE-Ig Protein Against Multiple SARS-CoV-2 Variants of Concern Demonstrated in K18-hACE2 Mouse and Syrian Golden Hamster Models": Suplement Figures S1 and S2

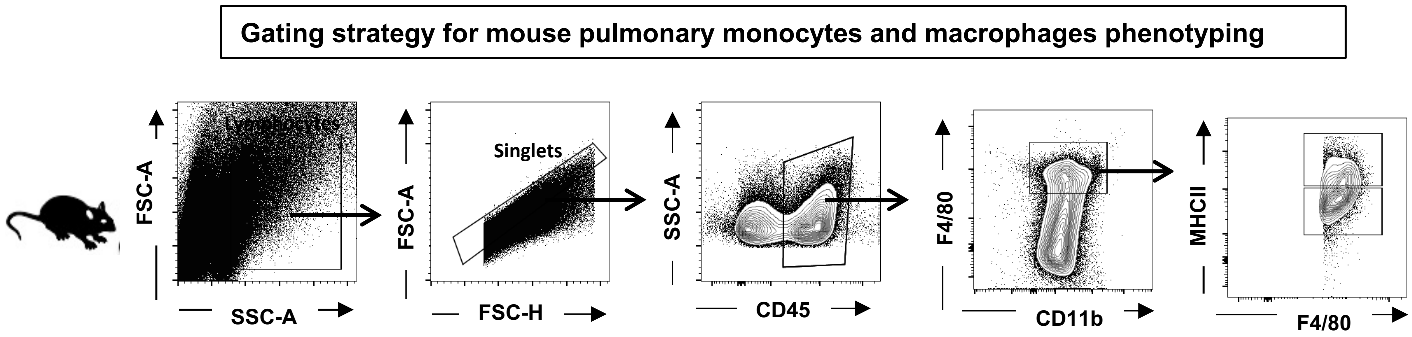


**Supplemental Figure S1**: **Gating strategy for mouse pulmonary monocytes and macrophages phenotyping**. Immune cells isolated from lungs of mice infected with SARS CoV2 were stained for macrophage population by flowcytometry. Live cells are gated using side-scatter (SSC) and forward-scatter (FSC), and doublets were excluded with FSC-A and FSC-H. F4/80$^+$CD11b$^+$CD45$^+$cells were then differentiated to monocyte or macrophage based on expression of MHCII.

**Kinetics of Type I, II & III interferon secretion in human lung epithelial cells (BEAS-2B) infected with SARS CoV-2/WA-2020**

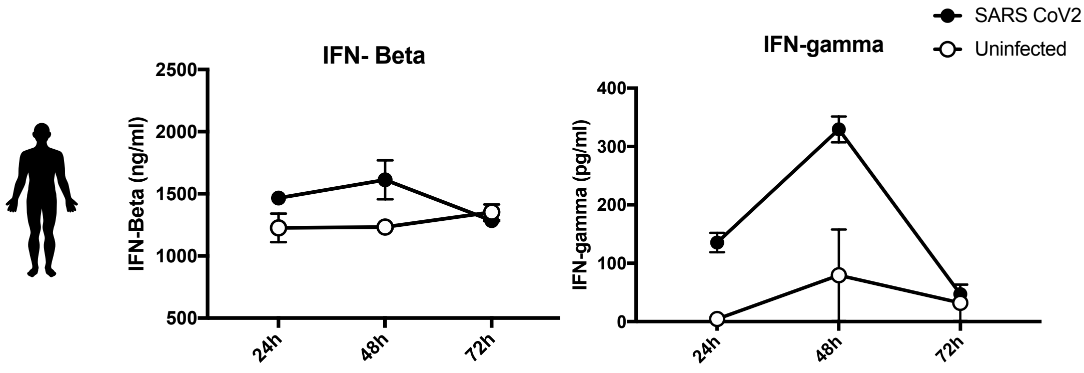


**Supplemental Figure S2**: Graph showing time-kinetics of IFN-β (left panel) and IFN-γ (right panel) released in supernatant of lung epithelial cells (BEAS-2B) infected with SARS CoV-2/WA-2020 at an MOI of 0.1, and estimated by ELISA.
